# Sequence similarity estimation by random subsequence sketching

**DOI:** 10.1101/2025.02.05.636706

**Authors:** Ke Chen, Vinamratha Pattar, Mingfu Shao

## Abstract

Sequence similarity estimation is essential for many bioinformatics tasks, including functional annotation, phylogenetic analysis, and overlap graph construction. Alignment-free methods aim to solve large-scale sequence similarity estimation by mapping sequences to more easily comparable features that can approximate edit distances efficiently. Substrings or *k*-mers, as the dominant choice of features, face an unavoidable compromise between sensitivity and specificity when selecting the proper *k*-value. Recently, subsequence-based features have shown improved performance, but they are computationally demanding, and determining the ideal subsequence length remains an intricate art. In this work, we introduce SubseqSketch, a novel alignment-free scheme that maps a sequence to an integer vector, where the entries correspond to dynamic, rather than fixed, lengths of random subsequences. The cosine similarity between these vectors exhibits a strong correlation with the edit similarity between the original sequences. Through experiments on benchmark datasets, we demonstrate that SubseqSketch is both efficient and effective across various alignment-free tasks, including nearest neighbor search and phylogenetic clustering. A C++ implementation of SubseqSketch is openly available at https://github.com/Shao-Group/SubseqSketch.

## 1 Introduction

Estimating the similarity between biological sequences is a fundamental task in bioinformatics, underpinning a wide range of applications including homology detection, gene annotation, and phylogenetic analysis. Traditionally, sequence similarity has been assessed with alignment-based methods, which attempt to find an optimal correspondence between characters from two or more sequences. While providing the most accurate results, these methods often suffer from high computational cost, especially when applied to large and divergent datasets.

Sketching-based methods have been developed to address this limitation. A sketch summarizes a long sequence into a small set of representative fingerprints that can be rapidly compared in place of the original sequences for similarity estimation. Together with its variants, the most widely used sketching method is MinHash (MH) [1]. In its simplest form, MH utilizes a hash function that maps each *k*-mer of a sequence to a number and only keeps the *k*-mer with the minimum hash value as the representative of that sequence. It is easy to see that the probability for two sequences to be represented by the same *k*-mer is proportional to the Jaccard similarity of the two sequences (viewed as sets of *k*-mers), namely, the number of shared *k*-mers between the sequences normalized by the total number of distinct *k*-mers among them. Hence, by repeatedly choosing min-*k*-mers with different hash functions and keeping track of the number of occurrences that the picked *k*-mers match between the two sequences, the Jaccard similarity can be estimated. In this process, the list of all representative *k*-mers of a sequence is called the MH sketch of this sequence. Two MH sketches are compared by the Hamming similarity – number of identical *k*-mers at the same indices. Order Min Hash (OMH) [19] extends this idea by estimating the weighted Jaccard similarity. Instead of picking one representative *k*-mer at a time, each entry of an OMH sketch is generated by picking several *k*-mers and putting them together following the original order in the sequence. OMH has been proved to be a locality-sensitive hashing family for the edit distance. A more comprehensive review of sketching algorithms for genomic data can be found in [21]. Note that both MH and OMH can be considered substring-based sketching methods because they pick substrings as the representatives. They therefore face the commonly observed difficulty in choosing a proper *k*: larger *k* is desirable to eliminate spurious matches but there are very few shared long *k*-mers even between closely related sequences.

To address this fundamental limitation of *k*-mers, several recent works [14, 11, 13] have advocated for the use of unrestricted subsequences instead. Subsequences relax the requirement that matching base pairs must be consecutive, allowing them to naturally tolerate gaps in the underlying – often unknown and computationally expensive – true alignment between sequences. This enables the identification of longer and hence more reliable matches, which in turn enhances the accuracy of downstream tasks. To fully leverage the benefits of subsequences, one must overcome a key algorithmic challenge: unlike the linear number of *k*-mers in a sequence, the number of subsequences grows exponentially, making MH-like strategies that rely on enumerating all candidates impractical. In this work, we seek to exploit structural properties of subsequences to overcome this computational barrier. To this end, we develop SubseqSketch, an efficient sketching method that summarizes long sequences into compact, subsequence-based features that are highly correlated with edit similarities. Through experiments on typical downstream applications, including nearest neighbor search and phylogenetic clustering, we demonstrate that SubseqSketch is both efficient and effective.

### 1.1 Related work

Recently, a sketching method named LexicHash [8] proposes to compare sketches based on the length of their common prefixes, rather than relying on fully matched *k*-mers. This has the effect of sketching with *k*-mers for all lengths *k* up to a predefined maximum value. However, LexicHash still suffers from the common issue of *k*-mer-based methods, namely, a small number of edits can destroy all long *k*-mer matches between two similar sequences. Furthermore, LexicHash is designed for the task of overlap detection, rather than estimating the similarity between two sequences. In particular, the authors define the LexicHash similarity score between two sequences as the length of the longest matching prefix among their sketches. So a score *k* only indicates that the two sequences share a common *k*-mer, which may be effective for detecting overlapping reads, but appears to be insufficient for edit similarity estimation (see Figure 4). In fact, choosing a proper distance function between sketches to facilitate a proper similarity estimation requires careful considerations for any sketching method, see Section 2.3 for further discussion.

To the best of our knowledge, the only existing subsequence-based sketching method is Tensor Slide Sketch (TSS) [11]. Instead of picking *k*-mers from the input sequence, TSS aims at producing a sketch by counting all subsequences. Since there is an exponential number of them, TSS has to group Subsequences in a smart way to facilitate counting. However, to make it efficient, TSS is restricted to count all short subsequences, which limits its capacity in distinguishing similar and dissimilar sequences.

## 2 SubseqSketch

The idea of SubseqSketch is to identify long common subsequences between input sequences through random sampling. Computing the sketch of a sequence *s* can be figuratively thought of as answering a survey in which each question asks whether *s* contains a randomly selected sequence as a subsequence. By comparing the answers of two sequences, their similarity can be estimated. We note that this idea does not work well with substrings (*k*-mers): As the number of *k*-mers in a sequence is negligible comparing to the number of length-*k* subsequences, the chance of successfully finding a reasonably sized common substring by random sampling is low, even between highly similar sequences. For a concrete example, according to Figure 3, if length-100 sequences are taking our survey, we can choose a query sequence to have length 25 and expect half of the answers to be “yes”. Furthermore, a matching “yes” answer for a pair of sequences suggests a (partial) alignment between them that involves at least a quarter of their bases. In contrast, if we were to ask whether a query 8-mer is a substring, the vast majority of sequences would answer “no”, resulting in a very weak, if functional at all, classifier for distinguishing between similar and dissimilar sequences.

While sampling long subsequences is beneficial for similarity estimation, it becomes computationally expensive on long inputs. In the following section, we introduce the concept of tokenization to effectively generalize the above strategy to genome-scale sequences. Combined with the idea of an “enhanced survey”, where binary yes/no questions are upgraded to integer-scale queries, we present the full-fledged SubseqSketch as an effective and efficient sketching method.

## 2.1 Tokenized subsequence

A sequence *x* of length *kt* over an alphabet Σ can be viewed as a sequence of *k* “tokens” each of which is a string of length *t*. We say *x* is a *tokenized subsequence* of a length-*n* sequence *s* if there is a list of indices 1 ≤ *i*_1_ *< i*_2_ *<* ··· *< i*_*k*_ ≤ *n* − *t* + 1 such that the length-*t* substring of *s* starting at *i*_*j*_ matches the *j*-th token of *x*. Note that when *t* = 1, a tokenized subsequence is a regular subsequence; it is not necessarily the case when *t >* 1, as the tokens are allowed to overlap, see Figure 1 for an example.

**Figure 1:**
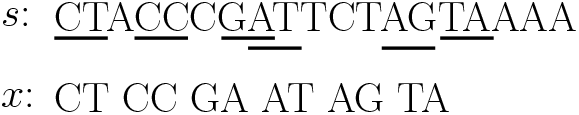
An example of tokenized subsequence. The bottom sequence *x* is tokenized with token size 2. It is a tokenized subsequence of the top sequence *s*, on which the corresponding tokens are underlined. Observe that *x* is not a regular subsequence of *s*.

### 2.2 Construction of SubseqSketch

To construct SubseqSketch for input sequences, we first generate a list *L* of random sequences of length *kt* each, where *k* and *t* are predefined parameters. We call *L* the list of testing subsequences. Two SubseqSketches are comparable only if they were generated with the same list *L*; in this sense, *L* serves as shared randomness in the sketching process, analogous to the shared random ordering of *k*-mers in MH sketches. Given an input sequence *s*, SubseqSketch takes a testing subsequence in *L* and determines the maximum number of its prefix length-*t* tokens that form a tokenized subsequence of *s*. The resulting vector consists of |*L*| integers, one for each testing subsequence. This vector is the sketch of *s*, denoted as SubseqSketch(*s*). See Figure 2 for an illustration.

**Figure 2:**
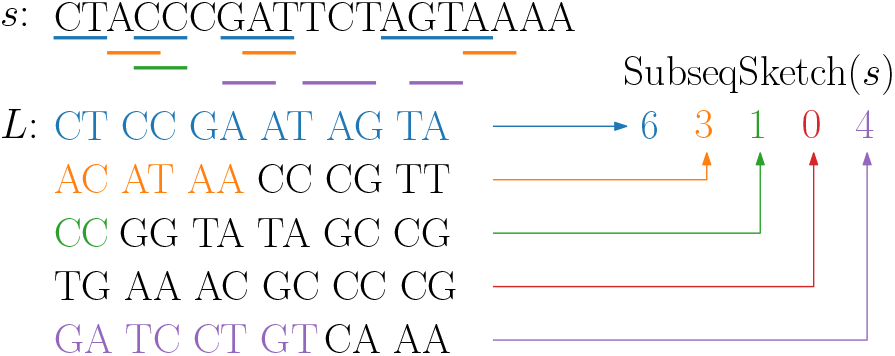
An illustration of SubseqSketch construction with *t* = 2, *k* = 6, and |*L*| = 5. For each testing subsequence in *L*, its maximum prefix tokens that form a tokenized subsequence of *s* are colored. Their matching tokens in *s* are underlined.

A straightforward linear scan computes the |*L*| sketch entries in *O*(|*L*| |*s*|) time. This worst-case time complexity can be improved by preprocessing the input sequence *s* to build an index that facilitates rapid lookup for the occurrence of the next token of a testing subsequence. For example, for token size *t* = 1, we can build an automaton on *s* in *O*(|*s*||Σ|) time and space. In the automaton, each character *s*_*i*_ stores |Σ| pointers. The pointer corresponds to *c* ∈ Σ points to the next appearance of *c* after *s*_*i*_ (or null if no *c* exists after *s*_*i*_). Then for a testing subsequence *x*, we can simply follow the pointers according to the characters of *x*, until either a null pointer is encountered or *x* is exhausted. This takes *O*(|*x*|) time for each testing subsequence so the total sketching time is *O*(|*s*| |Σ| + |*x*| |*L*|).

For larger token size, a similar idea can be applied: we can preprocess *s* to build a lookup table of size |Σ| ^*t*^ where each entry records the occurring positions of that token on *s*, either in a sorted array or some other data structures that supports quick search. Each testing subsequence can then be processed by following this lookup table until all tokens are used or the end of a position array is reached. This allows each integer in the sketch to be computed in *O*(*k* log |*s*|) time, instead of a *O*(|*s*|) linear search. We provide this preprocessing approach as an option in our implementation. However, through experiments we found that the linear search std::string::find provided in the standard C++ library is almost always faster. The overhead of preprocessing may only be justified for a large number of very long testing subsequences with a small token size, which is not a recommended setting for our sketching algorithm (see Section 2.4).

### 2.3 Choice of similarity function

The SubseqSketch of a sequence *s* provides a highly informative representation of *s*. To build intuition, consider two sequences *s* and *t*. If both sketches show large numbers at the same index, then *s* and *t* must share a long tokenized subsequence and hence likely similar in terms of the edit distance. Conversely, if one sketch has a large value while the other has a small value at the same index, it suggests that the sequences are likely dissimilar.

As with other sketching methods, a similarity measure over the sketches is required to translate the above intuition into a quantitative score that accurately reflects the true similarity between input sequences. Methods that compare sketches for equality at corresponding indices, such as MH and OMH, naturally employ Hamming similarity, which counts the number of matching entries between sketches. SubseqSketch, on the other hand, generates integer-valued vectors, enabling the use of a wide range of well-established distance/similarity metrics. We empirically evaluate a list of metrics using the data from Section 3.1. SubseqSketches are first computed, after which similarities scores are calculated using various metrics. The Pearson correlations between these scores and the ground truth edit similarities are reported in Table 1.

**Table 1:** The Pearson correlations between edit similarities and sketch similarity scores using various metrics.

| Metric | Pearson correlation |
| --- | --- |
| Canberra | 0.920 |
| Bray-Curtis | 0.919 |
| Correlation | 0.919 |
| Cosine | 0.918 |
| Hamming | 0.914 |
| Manhattan | 0.913 |
| Squared Euclidean | 0.901 |
| Jaccard | 0.881 |
| Euclidean | 0.857 |
| Minkowski | 0.809 |
| Chebyshev | 0.306 |

As shown in the table, cosine similarity is among the most effective metrics for producing estimates that are strongly correlated with the true edit similarity. According to its definition,

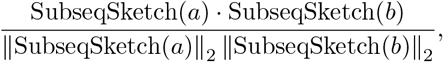

where denotes the vector dot product, pairwise cosine similarities between two sketching matrices can be computed using a single matrix multiplication (assuming the rows are normalized), which is highly optimized in modern hardware and numerical libraries. We therefore adopt cosine similarity between SubseqSketches in our implementation for its effectiveness and computational efficiency.

### 2.4 Choice of parameters

SubseqSketch has three parameters: the token size *t*, the number of tokens *k* in each testing subsequence, and the size |*L*| of the testing list. The parameter |*L*| controls the size of the sketches. In particular, a SubseqSketch takes |*L*| log *k* bits space to store. As with other sketching methods, increasing the sketch size improves estimation accuracy but comes at the cost of greater time and storage requirements. In the experimental sections, we compare the sketching methods at the same sketch size.

The parameters *t* and *k* are related. In the resulting sketches, each entry is an integer between 0 and *k*. If *t* is too large (for example, close to the input length *n*), most entries would be 0; on the other hand, if both *t* and *k* are small, most entries would max out at *k*, regardless of the input sequence *s*. Neither case is desirable as the sketches cannot provide a strong distinction between similar and dissimilar input sequences. Note that we can always choose a large *k* to ensure that few, if any, sketch entries reach the maximum value. However, this increases the sketch file size, as each entry requires log *k* bits – an inefficient use of space if most entries are significantly smaller than *k*.

We now try to derive an optimal choice of *k* for *t* = 1. In a recent paper [7], the authors motivated their sequence sampling method with an interesting puzzle (paraphrased): is the number of DNA 5-mers containing the substring ACGT the same as that for the substring AAAA? Astute readers will immediately answer “no” because it is impossible for a 5-mer to both start and end with ACGT – taking the union of the two disjoint groups gives the correct number – which is not the case for AAAA whose symmetry would cause the same strategy to double-count the 5-mer AAAAA.

As a curious extension, the same question can be asked, replacing substring with subsequence, namely, we do not require the containment to be consecutive. This seemingly more complicated version turns out to have a counterintuitively nicer answer: the number of *n*-mers containing a given *k*-mer as a subsequence is a function of *n* and *k*, independent of the choice of the *k*-mer. Consider a length-*k* sequence *x*, we count the number of length-*n* sequences *s* whose subsequence 1 ≤ *i*_1_ *< i*_2_ *<* · · · *< i*_*k*_ ≤ *n* is *x*. To avoid over-counting, we only count *s* if (*i*_1_, …, *i*_*k*_) is the first occurrence of *x* in *s*. It means the characters in *s* before *i*_1_ cannot be *x*_1_, leaving them |Σ| − 1 choices each. The same holds for regions in between *i*_*j*_ and *i*_*j*+1_, and finally all characters after *i*_*k*_ are free to be anything in Σ. This leads to 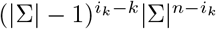 choices. Note that the expression only depends on *i*_*k*_ (i.e., any combination of *i*_1_, … *i*_*k*−1_ yields the same number), so we can group the terms and sum over choices of *i*_*k*_ to get the answer

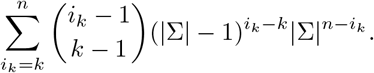

We emphasize that the calculation is independent of the chosen subsequence. An example is shown in Figure 3. We can then use the formula to compute a value of *k* such that at most a small threshold fraction (e.g., 0.01) of the sketch entries reach the maximum value *k*. In this example, *k* = 36 would suffice.

**Figure 3:**
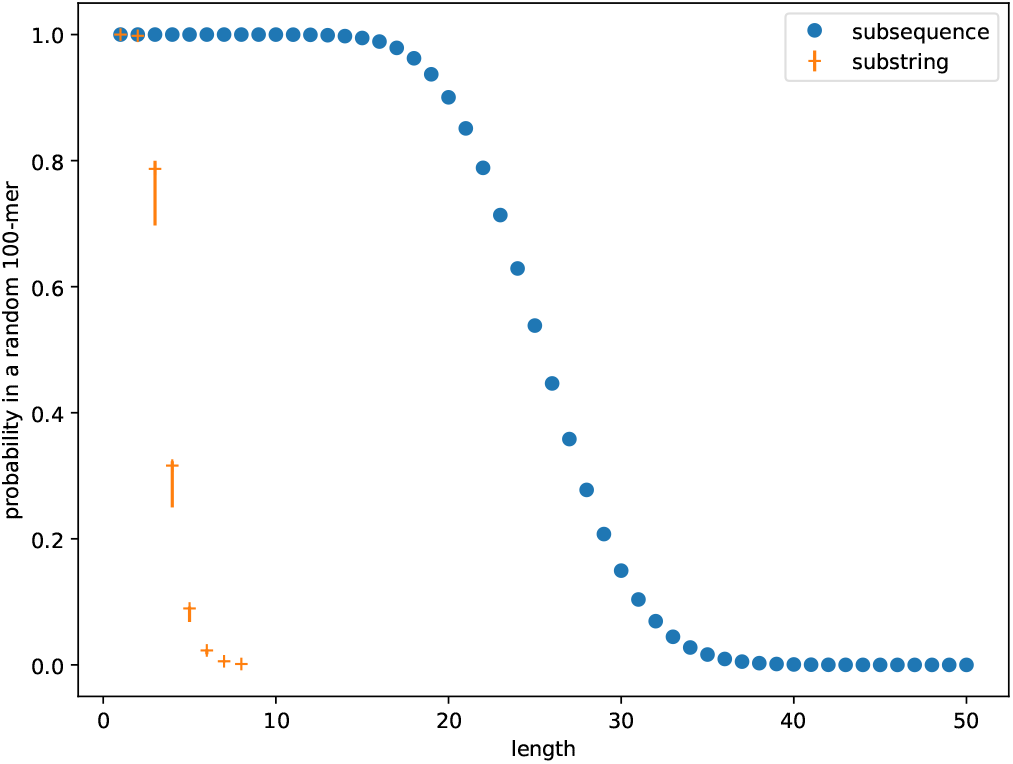
The fraction of length-100 sequences with |Σ| = 4 that contain a given subsequence (in blue) or substring (in orange) as a function of the length of the subsequence/substring. The blue dots are exact values computed according to the derived formula, they are the same regardless of the choice of the subsequence. The plots for substrings are empirical estimates; note that these can vary significantly across different *k*-mers, as indicated by the orange error bars for *k* up to 8.

**Figure 4:**
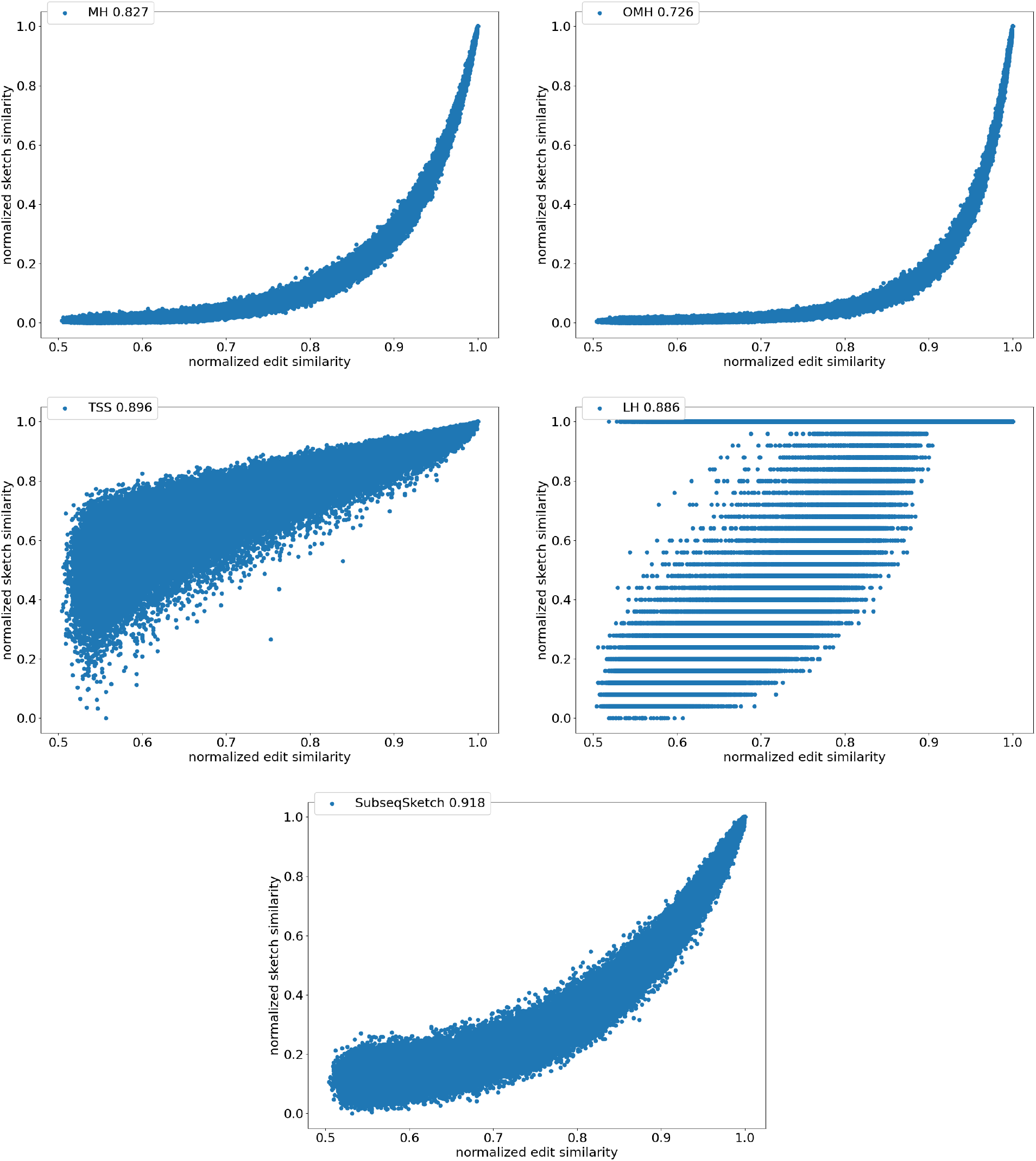
Correlation between normalized sketch similarities and normalized edit similarity on length *n* = 1000 sequences. The legend marks the name of the method and the Pearson correlation. All methods use sketch size 1000. Through parameter grid search, MH is configured to use *k*-mer size 8; OMH uses *k*-mer size 6 and *l* = 2; TSS uses *t* = 2, dimension 32, window size 0.1*n* = 100, stride size 0.01*n* = 10, as suggested in [11]. LexicHash uses maximum *k* 32. SubseqSketch uses token size 6.

For larger *t*, the derivation is not as neat. We can view a regular sequence *s* over the alphabet Σ as a tokenized sequence over the alphabet Σ^*t*^ and apply the above formula. But unlike adjacent characters in the original sequence, consecutive tokens with an overlap of length *t* − 1 are not independent, causing the formula to significantly overestimate. Since using a small *k* makes the sketching faster to compute and smaller to store, with an exception in Table 3, we fix *k* = 15 in the following experiments (namely, each entry in the sketch fits in 4 bits) and aim to choose *t* to ensure the sketching entries are neither too small nor maxed out. Table 2 provides empirical recommendations for *t* across common input sizes *n*.

**Table 2:** Empirical recommendations for parameter *t*.

| $n$ | $10^2$ | $10^3$ | $10^4$ | $10^5$ | $10^6$ | $10^7$ | $10^8$ | $10^9$ |
| --- | --- | --- | --- | --- | --- | --- | --- | --- |
| $t$ | 2 | 6 | 9 | 12 | 15 | 19 | 22 | 25 |

**Table 3:**
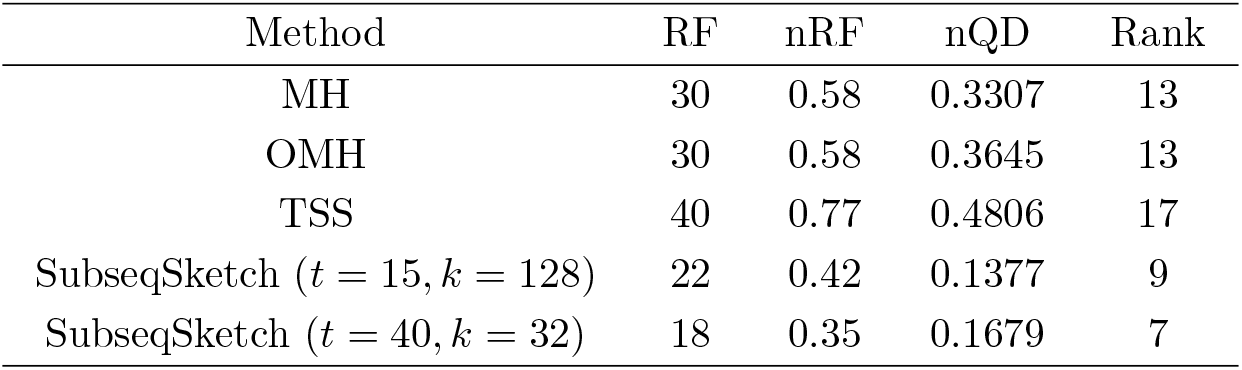
Phylogeny reconstruction results on 29 *E. coli* genomes. The RF, nRF, and nQD distances all measure topological disagreement between the reconstructed tree and the ground truth tree. A lower value indicates a more accurate reconstruction of the phylogeny. The Rank is based on the nRF distances among many tools tested by the AFproject. All methods use sketch size 10, 000. Through parameter grid search, MH is configured to use k-mer size 10 (in fact, multiple values of *k* between 10 and 30 all yield the same nRF distance, but *k* = 10 is slightly better on nQD); OMH uses k-mer size 22 and *l* = 3; TSS uses *t* = 5, dimension 100, window size 500, 000, stride size 100, 000. The parameters used by SubseqSketch are marked in parentheses.

### 2.5 Sample subsequences from input

Using randomly generated testing sequences is the best one can do in a data-oblivious setting, while better performance can usually be achieved if we can afford to adjust the sketches according to the input data. One idea to introduce data dependency is to sample subsequences from the input to form the testing list. This is particularly suitable when the input comprises a small number of sequences – for example, when estimating phylogenetic distances among a group of closely related genomes, as shown in Section 3.3. On the other hand, if the sketches are used to build an index of a large database of sequences to handle queries, it may not be practical to re-sketch the entire database with a new testing list for each query. In this situation, we simply use the data-oblivious version with a fixed list of randomly generated testing sequences and demonstrate in Sections 3.1 and 3.2 that it already achieves good performance.

## 3 Experiments

In this section, we first show a strong correlation between the cosine similarity of SubseqSketches with the edit similarity between simulated pairs of sequences. Then the sketch quality of SubseqSketch is tested on two sequence comparison tasks, the nearest neighbor search and phylogeny reconstruction. In each task, we compare SubseqSketch with competing methods on both simulated sequences and published benchmark datasets. For a fair comparison, each method is set to produce sketches of (roughly) the same size. A grid search is performed for each competing method to find the best parameters. Details are reported in each subsection.

### 3.1 Correlation between sketch similarity and edit similarity

To directly compare the sketch similarity against the desired but much more expensive to compute edit similarity, we generate 100, 000 random DNA sequences of length 1, 000. Each sequence is randomly mutated (an insertion, deletion, or substitution) for a random number of rounds up to 1, 000 to produce a pairing sequence. For each pair, we compute their exact edit similarity, as well as sketch similarities for SubseqS-ketch, MinHash (MH), Order Min Hash (OMH), Tensor Slide Sketch (TSS) and LexicHash (LH). For each sketching method the Pearson correlation between the exact edit similarity and the sketch similarity over the 100,000 pairs of sequences is reported. MH, OMH, and TSS use the implementation of [11]. LH uses the implementation of [8].

Figure 4 shows the scatter plots of all the pairs under different sketching methods. The horizontal axis marks the normalized edit similarity which is computed as one minus the edit distance divided by sequence length. The vertical axis shows the sketch similarities which are normalized to the range [0, 1]. Observe that SubseqSketch achieves the best Pearson correlation. Both MH and OMH are good estimators for sequences with high edit similarities but struggle to distinguish dissimilar sequences with edit similarity between 0.5 and 0.8. The TSS and LH similarities show a visually more linear relationship with the edit similarity and consequently exhibit higher Pearson correlations than MH and OMH. But they both suffer from extremely large variance, especially for dissimilar sequences, which makes it difficult to interpret their estimation in practical applications. SubseqSketch strikes a balance between the ability to estimate the full range of edit similarity and the estimation variance.

As with other sketching methods, the variance of SubseqSketch can be reduced by using a larger sketch. For all the experiments, we measure the size of a sketch as the number of entries in it (sometimes called its dimension), and all methods are configured to produce the same number of entries (except for TSS, which we follow the suggestion in [11] even though it produces a larger sketch). However, in real applications, the actual space needed to store the sketches is a more relevant measure. Recall that each entry of SubseqSketch can be stored in 4 bits (ref. Section 2.4) which is four times smaller than an entry of MH (16 bits for *k* = 8), six times smaller than OMH (24 bits for *k* = 6 and *f* = 2), and eight times smaller than TSS and LH (32-bit float/int). Thus, given a fixed amount of disk space, SubseqSketch can utilize more testing subsequences than the number of *k*-mers MH or OMH can select, thereby achieving a similar or better variance. In the experiments, we do not exploit this practical advantage, opting instead to use the same number of sketch entries across all methods.

### 3.2 Nearest neighbor search

The task of nearest neighbor search asks to find the top-*T* most similar sequences for a query among a large database. Since computing the exact edit distance between the query and every sequence in the database is computationally prohibitive, a common approach is to map database sequences into a well-studied metric space where efficient nearest neighbor indexing is readily available (for example, the hierarchical navigable small world index [17]). A query can then be mapped into the same space, and the nearest neighbors according to the index are reported as approximations of the true nearest neighbors in the original sequence space. In this experiment, we choose to not include any indexing because the accuracy of the index may affect the final results. Following the pipeline of CNN-ED [4], a tool that performs sequence nearest neighbor search using a learned embedding for edit distance, we compute the sketch distances between a query and all sequences in the database and report the top-*T* nearest neighbors. It is worth noting that computing sketch distances is much more scalable than computing edit distances.

We show results on two widely used datasets GEN50kS and GEN20kL from [26] which are also bench-marked in the CNN-ED paper. The GEN50kS dataset contains 50, 000 sequences with an average length 5, 000. The GEN20kL dataset contains 20, 000 sequences with an average length 20, 000. The CNN-ED pipeline splits each dataset into three disjoint sets: a training set with 1, 000 sequences, a query set with 1, 000 sequences, and a base set containing the remaining sequences. It then computes the all-vs-all edit distances between the query set and the base set to form the ground truth for the nearest neighbor search. For the sketching methods, the training set is not used.

To evaluate the performance of different methods, we plot the commonly used recall-item curves in Figure 5 and Figure 6. For a figure labeled top-*T*, the *T* nearest neighbors of a query in the base set according to the edit distances are considered true neighbors. The horizontal axis represents the number of neighbors (items) each method is allowed to report (according to their respective sketch/embedding distances) and the vertical axis marks the fraction of true neighbors being reported (recall). The CNN-ED pipeline presents full-range results – from reporting a single item to reporting all items – which, while not practical for typical use cases (where only the top-*T* neighbors are retrieved), allows for plotting complete performance curves.

**Figure 5:**
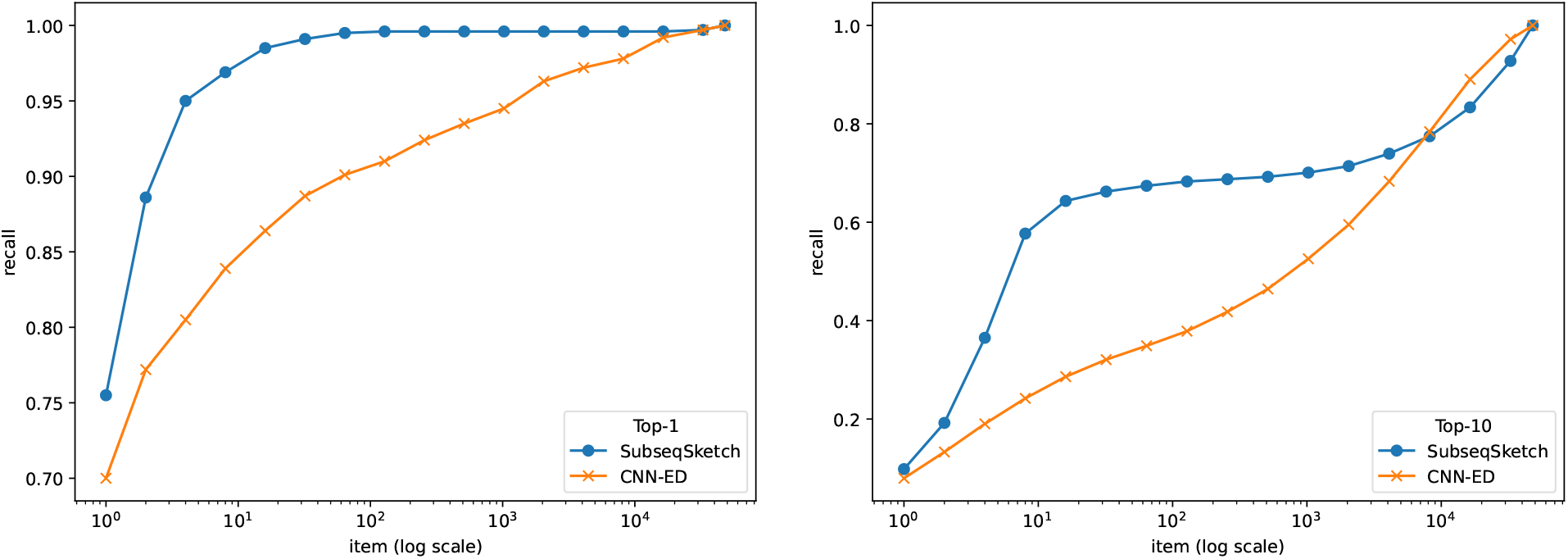
Recall-item curves of different methods on the GEN50kS dataset. All methods output vectors of dimension 200. SubseqSketch uses token size 6. Left: ground truth is the top-1 nearest neighbor by edit distance. Right: ground truth contains the top-10 nearest neighbors by edit distance.

**Figure 6:**
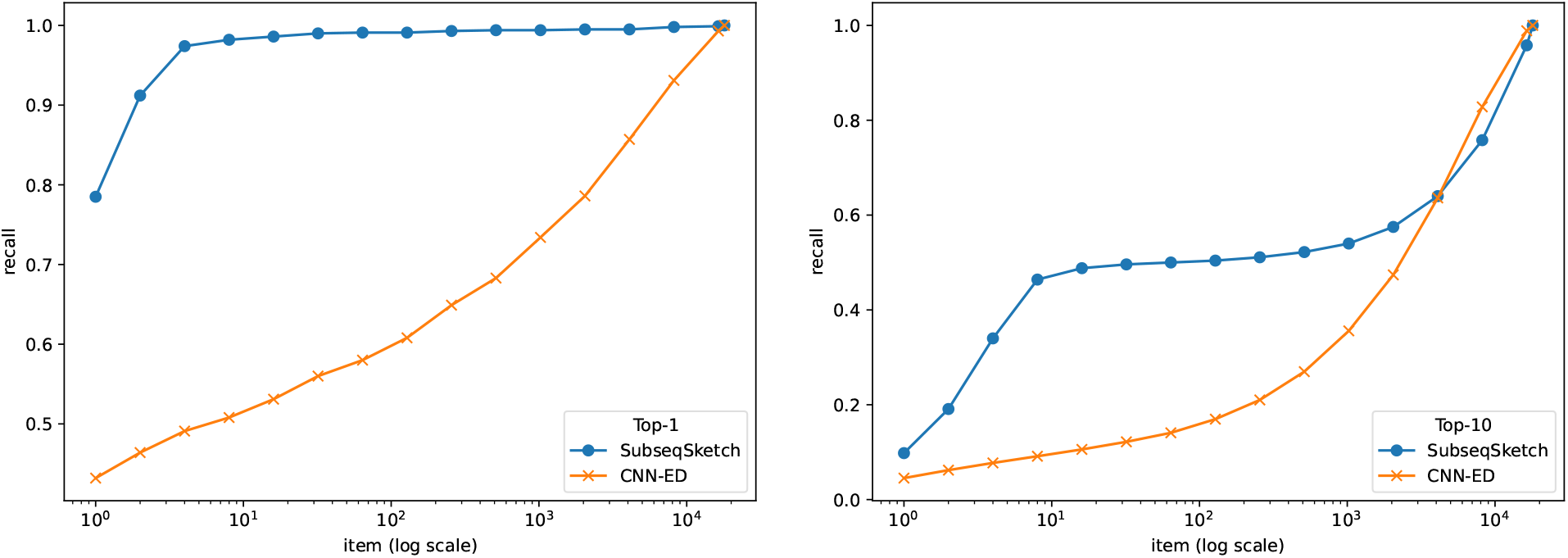
Recall-item curves of different methods on the GEN20kL dataset. All methods output vectors of dimension 128. SubseqSketch uses token size 7. Left: ground truth is the top-1 nearest neighbor by edit distance. Right: ground truth contains the top-10 nearest neighbors by edit distance.

In this experiment we restrict our comparison to CNN-ED, which was shown to outperform other non-machine learning methods such as the CGK embedding [3]. The CNN-ED results are obtained by the implementation of [4]. It is a deep convolutional neural network model which we trained for 50 epochs following the reported hyperparameters in the original paper. For a fair comparison, SubseqSketch is configured to produce vectors of the same length as the embedding dimensions of CNN-ED. Observe that SubseqSketch consistently outperforms CNN-ED by a large margin. This is a surprising result. It is commonly believed (which is often, though not always, justified) that machine learning models can outperform traditional algorithmic methods because the models can learn data-dependent features that the data-oblivious algorithms cannot take advantage of. In [4], the CGK embedding [3] was shown to produce a worse result than CNN-ED on this task, even though it is an edit distance embedding with theoretical guarantees. Our result here demonstrates that there is a gap between theoretical bounds and practical performance which warrants further investigation. In particular, we conjecture that SubseqSketch can also provide some guarantees on the distortion as a randomized embedding function for the edit distance, though a theoretical proof seems difficult.

### 3.3 Phylogeny reconstruction

Phylogeny reconstruction is another common task that can be used to evaluate the performance of alignment-free methods. Given a set of biologically related genomes, the goal is to build a phylogeny on them based on pairwise similarities/distances estimated by the sketches. The result can then be compared with a ground truth tree constructed from some biological model or multiple sequence alignment. We test on two datasets for this task: one is a simulation of a simple mutation model similar to that used in [19]; the other is a set of 29 assembled *E. coli* genome sequences collected in [25].

For both datasets, an all-vs-all distance matrix is computed for each method. For the simulated dataset, the matrices are used to build the phylogenies with the neighbor-joining algorithm implemented in the biotite package [12]. The normalized Robinson-Foulds (nRF) distances between the constructed trees and the ground truth tree are then calculated with the ETE toolkit [9]. The nRF distance measures the dissimilarity of branching patterns between two trees and ignores branch lengths. A value of 0 means the two phylogenies have the identical tree topology, whereas a value of 1 indicates the two trees are maximally dissimilar. For the real *E. coli* genome sequences, the AFproject [27] (a benchmark project for alignment-free sequence analysis tools) provides a web interface where the phylogenies can be computed from the uploaded distance matrices. The nRF distances are then reported by comparing the resulting trees against a ground truth tree built from multiple sequence alignment. It also provides the normalized Quartet Distance (nQD) as an additional measure for topological disagreement. On the website, many alignment-free phylogeny reconstruction tools are ranked based on the nRF distances achieved.

Following the experiment in [19], we simulate a family of sequences using a simple mutation model that includes both point mutations and mobile genomic elements, commonly found in bacterial genome rearrangements, known as insertion sequences (IS). The simulated sequences form a perfect binary tree. The root of the tree is a random sequence of length 10, 000; it is considered as the 0-th generation genome. To obtain the *i*-th generation, each sequence in the (*i* − 1)-th generation produces two children genomes by independent and random point mutations with mutation rate 0.01%. Then a random IS of length 500 is inserted at a random position for each newly generated *i*-th generation genomes. Note that the IS is shared among all sequences in the same generation, but the inserting positions can be different. See Figure 7 for an illustration. Although simple and somewhat unrealistic, this model produces a solid ground truth phylogeny and allows us to investigate the effectiveness of different sketching methods to recover the mixed history of point mutations and large insertion events.

**Figure 7:**
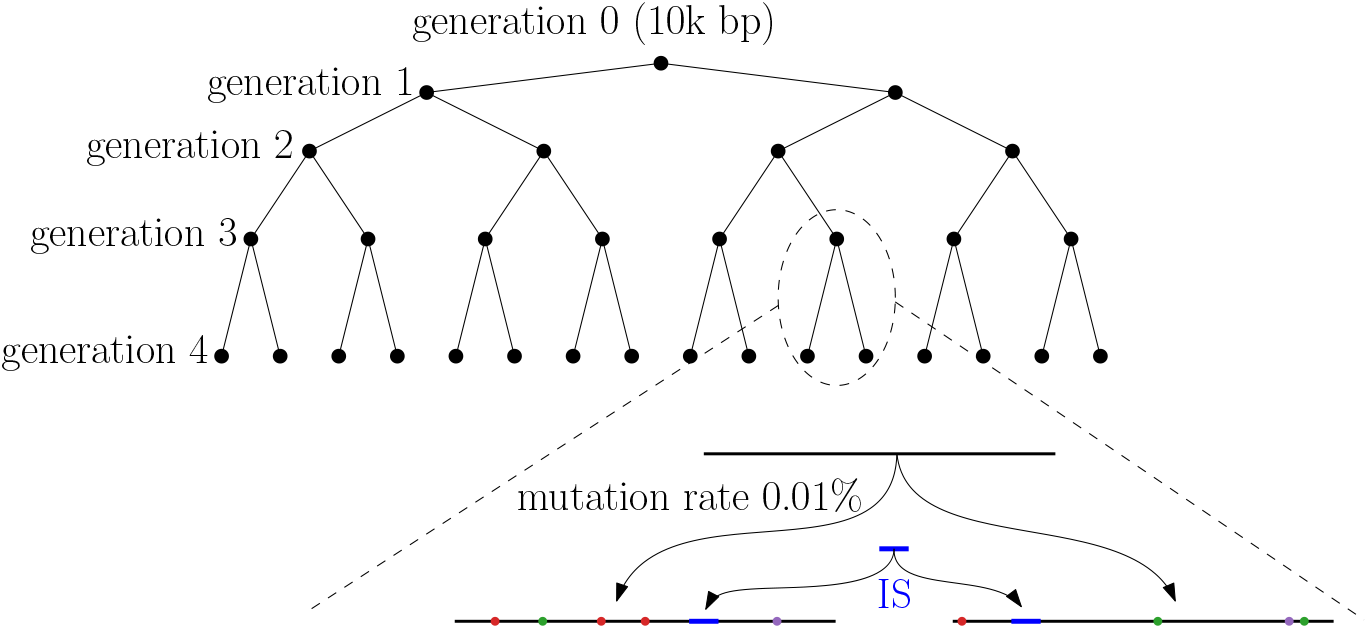
An illustration of the simulated phylogeny. In the zoomed-in view at the bottom, the top segment represents a sequence from the 3-rd generation. Its two children in the 4-th generation are obtained by random point mutations represented by colored dots. The blue segment represents the common IS inserted into each sequence in the 4-th generation.

Figure 8 shows the nRF distances achieved by each method on progressively larger inputs from the simulated dataset. The horizontal label *i* means all the 2^*i*^ sequences from the *i*-th generation are used as input sequences. Not surprisingly, pairwise edit distance (ED) most accurately captures the mutation history, at the cost of significantly longer computation time (see Figure 9). Among the sketching methods, SubseqSketch constructs the best phylogeny for generation 6 and larger inputs. Furthermore, the nRF distances obtained by SubseqSketch exhibits a strong correlation with those achieved by the exact edit distances, indicating it can be used as a faithful approximation of the expensive edit calculation. In contrast, although MH and OMH produce trees with smaller nRF distances for the smaller input sets, they both show some inverse relation with the nRF using edit distances (from generation 3 to 4, the nRF distances of trees constructed by edit distance increased, but the nRF distances for MH decreased; similarly from generation 4 to 5 for OMH). LH is omitted from this experiment because its implementation choice for boundary handling tends to assign the maximum similarity score to pairs sharing a short matching suffix (see the line at normalized similarity score 1 in Figure 4). While this may be appropriate for the overlap detection task that LH is designed for, it hinders accurate phylogeny reconstruction on our datasets.

**Figure 8:**
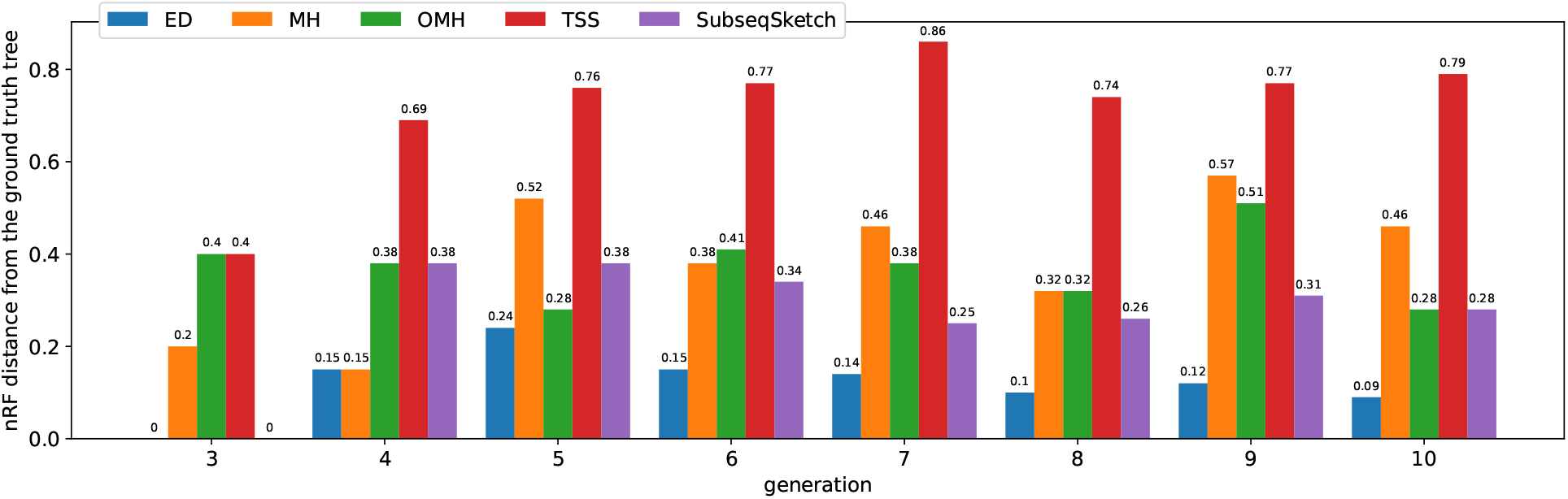
Normalized RF distances achieved by each method on the simulated dataset. A lower nRF distance indicates the constructed phylogeny is more similar to the ground truth tree. All methods use sketch size 256. Through parameter grid search, MH is configured to use *k*-mer size 8; OMH uses *k*-mer size 6 and *l* = 2; TSS uses *t* = 4, dimension 16, window size 1, 000, and stride size 100. SubseqSketch uses token size 5.

**Figure 9:**
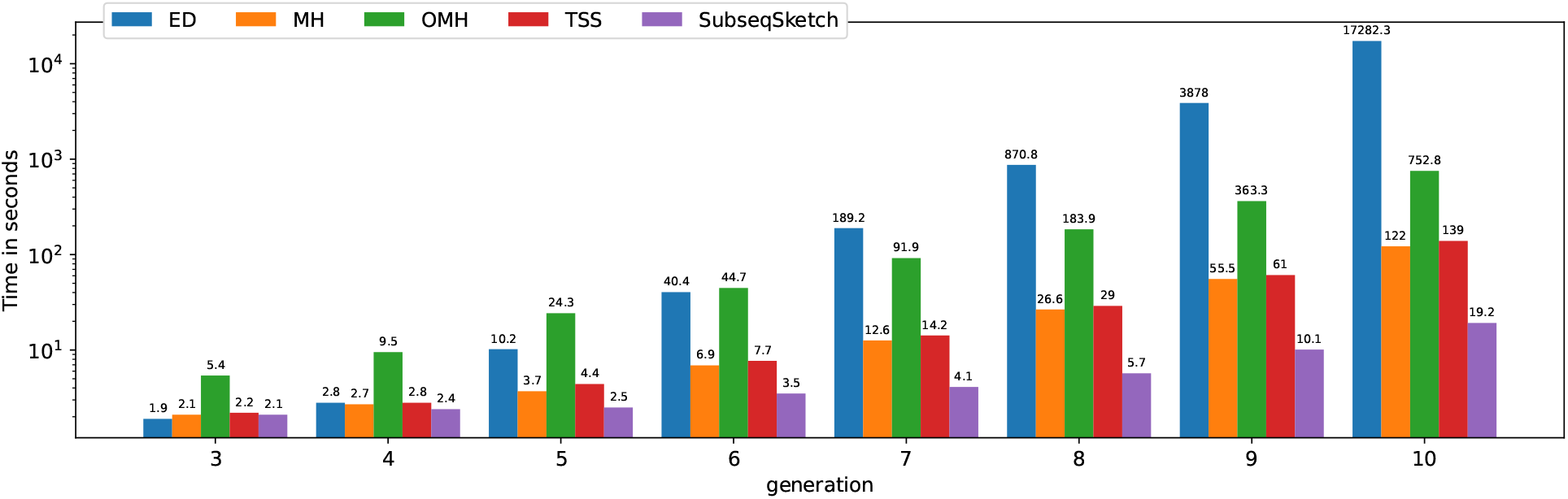
Time spent by each method in seconds (log scale). All experiments run on a server with an Intel(R) Xeon(R) Gold 6148 CPU @ 2.40GHz. Edit distance is computed with the Python package Levenshtein. MH, OMH, and TSS are computed using the implementation of [11].

We also plot the running time of each sketching method in Figure 9 to demonstrate the efficiency of SubseqSketch. As expected, all the sketching methods are much faster than computing the all-vs-all exact edit distances. Among them, SubseqSketch is consistently the fastest, regardless of the number of input sequences. More specifically, SubseqSketch achieves a 6× speedup compared to the second fastest method (MH).

Results for the real *E. coli* dataset are summarized in Table 3. On the AFproject website, nearly 100 tools (include different configurations for the same tool) are ranked based on the nRF distance. SubseqSketch is ranked 7th and there are 12 tools that achieve smaller nRF distances due to ties. It is worth pointing out that the higher ranked ones are tools designed specifically for the task of phylogeny reconstruction, which are often based on some sketching method but also apply biological and algorithmic heuristics to adjust the sketch distance matrix. Since SubseqSketch is a sketching method rather than a complete tool for phylogeny, here we aim to evaluate the sketch quality without those adjustments. By using the raw distance matrices, SubseqSketch constructs the best phylogeny (closest to the ground truth) among MH, OMH, and TSS.

In this task, since there are only 29 genomes, we can afford to sample the testing subsequences from the input to further improve the quality of SubseqSketch. Because the inputs are all closely related, this sampling strategy also enables us to use a much larger token size *t* = 40 to achieve an even better result than the recommended *t* = 15. From Table 3, it is evident that setting *t* = 40 significantly improves accuracy.

## 4 Discussion

We presented SubseqSketch, a subsequence-based sketching method that is both effective and efficient at sequence similarity estimation. Comparing to the widely used MH, OMH, TSS, and LH sketches, SubseqSketch requires smaller space, is faster to compute, and achieves a stronger correlation with the edit similarity. It delivers strong performance in two alignment-free tasks: nearest neighbor search and phylogeny reconstruction. In particular, it outperforms a machine learning edit distance embedding model by a large margin which suggests our method indeed captures critical features of the sequences being sketched.

A large body of work that we intentionally excluded from our experiments consists of seeding-based methods. The simplest seeds are *k*-mers, representing fixed-length consecutive exact matches in the sequences. More advanced *k*-mer selection schemes exist, such as minimizer [24, 18], syncmer [5] and *k*-min-mer [6]. Seeds sampled from subsequences, either with limited patterns such as spaced seed [2, 15] and strobemer [22, 16, 23], or fully unrestricted such as SubseqHash [14, 13], have been shown to deliver better performance but are usually more expensive to compute. While both sketching and seeding utilize some common techniques, for example, the minimizer seeds are obtained by applying MH [1] on each window, they differ significantly in their goals, representations and usage. Seeding methods aim to identify local regions of similarity between sequences, providing fine-grained information about where and how sequences resemble each other. This often comes at the cost of increased memory footprint and computational overhead. Specifically, seeding methods typically extract seeds from a relatively small sliding window over a longer input sequence. By generating one or more seeds from each overlapping window^1^, the number of seeds for a sequence of length *n* is usually Θ(*n*). In contrast, sketching methods prioritize efficiency by transforming sequences into compact, low-dimensional representations that enable fast, global similarity estimation. For example, an *E. coli* genome with several million base pairs is condensed to a length 10, 000 vector by each sketching method in the above experiment. Unlike seeds, which are often used temporarily during computation and then discarded, sketches are typically stored and reused, serving as compact indices in databases containing vast numbers of sequences.

There are numerous interesting directions that call for further investigations. From the theoretical perspective, a deeper understanding of SubseqSketch, and subsequence-based features in general, can be beneficial for better algorithmic designs as well as guiding practical applications. Many methods compared in the experiments come with theoretical guarantees: MH is an unbiased estimator for the Jaccard similarity; OMH is a locality-sensitive hashing (LSH) family for the edit distance; and CGK is an embedding for the edit distance with a quadratic distortion. Given the superior performance of SubseqSketch against these methods, it is natural to consider what bounds can be proved on it. More specifically, we are curious if SubseqSketch is an LSH, and if so, does it offer better hash collision probabilities? Or is it an embedding with provable small distortion for the edit distance? In that case, study the relation between its parameters and the achieved distortion can help to make informed decisions in practical use.

On the application side, there are several potential approaches to enhance SubseqSketch. For example, Mash [20] is a popular tool for genome distance estimation. It is based on MH whose estimation does not exhibit the strongest correlation with edit distance. However, by applying a simple Poisson model to adjust the MH score, Mash produces a distance that closely approximates the mutation rate on real datasets. Since SubseqSketch starts with a more accurate estimation, it is reasonable to believe that similar techniques can be applied to further improve its performance.

A related question concerns the similarity function used by SubseqSketch. The cosine similarity was chosen for its effectiveness and simplicity. While it matches our intuition that sketches of similar sequences should have near identical corresponding entries and therefore should be roughly pointing to the same direction in the sketch vector space, the cosine similarity explicitly ignores the magnitude of the vectors. In the extreme case, a sketch full of 1’s is considered to have the maximum similarity with another sketch full of 10’s. This greatly diverges from the designed meaning of the SubseqSketch entries – the first sequence barely contains any testing subsequences whereas the second contains large portions of each testing subsequence – they must be very different! Exploring different similarity functions that can better incorporate the expected interpretation of the entries can therefore potentially make SubseqSketch more accurate.

Yet another observation is that SubseqSketch is sensitive for globally well-aligned sequences but can struggle with ones that only share meaningful local alignments. For example, we cannot expect a genome comprising millions of base pairs to produce a SubseqSketch similar to that of a 100-base-pair short read.

Other sketching methods such as MH also suffer from these situations and special variants such as FracMin-Hash [10] are designed to handle them differently. As another example, in building overlap graphs for genome assembly, one needs to identify overlapping pairs of sequences that contain additional unaligned prefixes and suffixes. Suppose that the tail of sequence *a* overlaps with the head of sequence *b*. Since SubseqSketch tests for subsequences from left to right and stops immediately when the next token cannot be found, the sketches will be disproportionally skewed: because *b* does not have the beginning part of *a*, testing subsequences fully live inside *a* can produce 0’s for *b*, even if *b* contains long suffixes of them. We hope to see diverse adaptations of SubseqSketch designed to address these various challenges.

## Acknowledgments

This work is supported by the US National Science Foundation (2145171 to M.S.) and by the US National Institutes of Health (R01HG011065 to M.S.).

## Footnotes

1 There also exist seeding schemes without a window guarantee, such as syncmer [5].

## References

[1] Andrei Z Broder. On the resemblance and containment of documents. In Proceedings. Compression and Complexity of SEQUENCES 1997 (Cat. No. 97TB100171), pages 21–29. IEEE, 1997.

[2] Andrea Califano and Isidore Rigoutsos. FLASH: A fast look-up algorithm for string homology. In Proceedings of IEEE Conference on Computer Vision and Pattern Recognition (CVPR’93), pages 353–359. IEEE, 1993.

[3] Diptarka Chakraborty, Elazar Goldenberg, and Michal Koucký. Streaming algorithms for embedding and computing edit distance in the low distance regime. In Proceedings of the 48th ACM Symposium on Theory of Computing (STOC’16), pages 712–725, 2016.

[4] Xinyan Dai, Xiao Yan, Kaiwen Zhou, Yuxuan Wang, Han Yang, and James Cheng. Convolutional embedding for edit distance. In Proceedings of the 43rd international ACM SIGIR conference on Research and Development in information retrieval, pages 599–608, 2020.

[5] Robert Edgar. Syncmers are more sensitive than minimizers for selecting conserved k-mers in biological sequences. PeerJ, 9:e10805, 2021.

[6] BariŞ Ekim, Bonnie Berger, and Rayan Chikhi. Minimizer-space de Bruijn graphs: Whole-genome assembly of long reads in minutes on a personal computer. Cell Systems, 12(10):958–968, 2021.

[7] Martin C Frith, Jim Shaw, and John L Spouge. How to optimally sample a sequence for rapid analysis. Bioinformatics, 39(2):btad057, 2023.

[8] Grant Greenberg, Aditya Narayan Ravi, and Ilan Shomorony. Lexichash: sequence similarity estimation via lexicographic comparison of hashes. Bioinformatics, 39(11):btad652, 10 2023.

[9] Jaime Huerta-Cepas, François Serra, and Peer Bork. Ete 3: Reconstruction, analysis, and visualization of phylogenomic data. Molecular Biology and Evolution, 33(6):1635–1638, 02 2016.

[10] Luiz Irber, Phillip T Brooks, Taylor Reiter, N Tessa Pierce-Ward, Mahmudur Rahman Hera, David Koslicki, and C Titus Brown. Lightweight compositional analysis of metagenomes with fracminhash and minimum metagenome covers. BioRxiv, pages 2022–01, 2022.

[11] Amir Joudaki, Gunnar Ratsch, and André Kahles. Fast alignment-free similarity estimation by tensor sketching. bioRxiv, 2020.

[12] Patrick Kunzmann, Tom David Müller, Maximilian Greil, Jan Hendrik Krumbach, Jacob Marcel Anter, Daniel Bauer, Faisal Islam, and Kay Hamacher. Biotite: new tools for a versatile python bioinformatics library. BMC bioinformatics, 24(1):236, 2023.

[13] Xiang Li, Ke Chen, and Mingfu Shao. Efficient seeding for error-prone sequences with subseqhash2. bioRxiv, pages 2024–05, 2024.

[14] Xiang Li, Qian Shi, Ke Chen, and Mingfu Shao. Seeding with minimized subsequence. Bioinformatics, 39(Supplement 1):i232–i241, 06 2023.

[15] Bin Ma, John Tromp, and Ming Li. Patternhunter: faster and more sensitive homology search. Bioinformatics, 18(3):440–445, 2002.

[16] Benjamin Dominik Maier and Kristoffer Sahlin. Entropy predicts fuzzy-seed sensitivity. bioRxiv, page 2022.10.13.512198, 2022.

[17] Yu A Malkov and Dmitry A Yashunin. Efficient and robust approximate nearest neighbor search using hierarchical navigable small world graphs. IEEE transactions on pattern analysis and machine intelligence, 42(4):824–836, 2018.

[18] Guillaume Marçais, Dan DeBlasio, and Carl Kingsford. Asymptotically optimal minimizers schemes. Bioinformatics, 34(13):i13–i22, 2018.

[19] Guillaume Marçais, Dan DeBlasio, Prashant Pandey, and Carl Kingsford. Locality-sensitive hashing for the edit distance. Bioinformatics, 35(14):i127–i135, 2019.

[20] Brian D Ondov, Todd J Treangen, Páll Melsted, Adam B Mallonee, Nicholas H Bergman, Sergey Koren, and Adam M Phillippy. Mash: fast genome and metagenome distance estimation using minhash. Genome biology, 17:1–14, 2016.

[21] Will PM Rowe. When the levee breaks: a practical guide to sketching algorithms for processing the flood of genomic data. Genome biology, 20:1–12, 2019.

[22] Kristoffer Sahlin. Effective sequence similarity detection with strobemers. Genome Research, 31(11):2080–2094, 2021.

[23] Kristoffer Sahlin. Strobealign: flexible seed size enables ultra-fast and accurate read alignment. Genome Biology, 23(1):1–27, 2022.

[24] Saul Schleimer, Daniel S Wilkerson, and Alex Aiken. Winnowing: local algorithms for document fingerprinting. In Proceedings of the 2003 ACM SIGMOD International Conference on Management of Data (SIGMOD/PODS’03), pages 76–85, 2003.

[25] Huiguang Yi and Li Jin. Co-phylog: an assembly-free phylogenomic approach for closely related organisms. Nucleic Acids Research, 41(7):e75–e75, 01 2013.

[26] Haoyu Zhang and Qin Zhang. Embedjoin: Efficient edit similarity joins via embeddings. In Proceedings of the 23rd ACM SIGKDD international conference on knowledge discovery and data mining, pages 585–594, 2017.

[27] Andrzej Zielezinski, Hani Z Girgis, Guillaume Bernard, Chris-Andre Leimeister, Kujin Tang, Thomas Dencker, Anna Katharina Lau, Sophie Röhling, Jae Jin Choi, Michael S Waterman, et al. Benchmarking of alignment-free sequence comparison methods. Genome biology, 20:1–18, 2019.

